# Standing Genomic Variation inferred by popGWAS Predicts Seedling Drought Survival Experiment in *Fagus sylvatica*

**DOI:** 10.64898/2026.07.13.738179

**Authors:** Linda Eberhardt, Friederike Reuss, Markus Pfenninger

## Abstract

Intensifying drought regimes across Central Europe, driven by global climate change, pose an increasing threat to forest regeneration, with the seedling stage representing the primary demographic bottleneck for long-lived tree species such as *Fagus sylvatica*. While intraspecific phenotypic variation in drought resistance among beech seedlings has been documented, the genomic basis of this variation remains poorly understood. Existing provenance trial approaches capture broad-sense heritability but lack the resolution to identify specific causal loci. To address this gap, we conducted a controlled soil drought experiment with *Fagus sylvatica* seedlings and applied a population-level genome-wide association study (GWAS), and leveraged individual survival length as a direct fitness proxy. We detected substantial variation in survival duration (10–40 days) among populations, with corresponding allele frequency differences at candidate loci whose functional annotations partially reflect known responses to environmental stress. Notably, seedlings sourced from an official seed bank consistently underperformed relative to wild-collected material, suggesting that current seed sourcing practices may inadvertently deplete adaptive genetic diversity. Our results demonstrate that standing genomic variation in natural *Fagus sylvatica* populations predicts differential seedling drought survival, and that the species retains substantial genetic potential to cope with prolonged spring drought when natural genetic diversity is preserved. These findings have direct implications for assisted migration strategies and seed sourcing guidelines under projected climate scenarios.

## 1. Introduction

Global climate change is driving drought events increasing in frequency, intensity and duration (Gazol et al., 2025), with severe drought periods in 2018 and 2019 triggering large-scale tree mortality across Central Europe (Schiefer et al., 2025). Historical records spanning 180 years indicate that summer droughts have intensified while winter droughts have declined (Neimry et al., 2025), and in addition, the frequency of spring droughts has risen sharply since 2003(Haslinger & Mayer, 2023). Spring droughts are of particular ecological significance for the establishment of forest trees of younger age classes, as they expose seedlings and young growth to increased soil desiccation and atmospheric drought stress before their root systems and the protective canopy closure are fully established (Lazarus et al., 2018) . This increasing phenological mismatch between evolved moisture expectations and shifting precipitation regimes constitutes a severe physiological stressor for early-stage vegetation. Young trees are structurally ill-equipped to cope with such conditions: their shallow root systems are confined to surface soil layers most susceptible to rapid drying, and their limited secondary growth provides little protection against desiccative water loss through above-ground tissues (Padilla & Pugnaire, 2007). Consequently, even brief drought episodes that cause surface soil drying disproportionately impact seedlings and young growth relative to established trees (Príncipe, 2019). While intraspecific variation in drought response among *Fagus sylvatica* seedlings has been documented at the phenotypic level (Aranda et al., 2015; Bolte et al., 2016; Cocozza et al., 2016; Kurath et al., 2025; Rose et al., 2009; Tognetti et al., 1995; Wang et al., 2021), the genomic basis of this variation remains poorly understood. Existing studies rely predominantly on provenance trial designs that capture broad-sense heritability but lack the resolution to identify specific genomic loci contributing to drought survival. This gap is particularly consequential for the seedling stage, which represents the primary demographic bottleneck for forest regeneration (Gauzere et al., 2016), yet has received comparatively little attention in genomic adaptation studies of European beech. Identifying the genomic loci that confer resilience at this stage of tree growth is therefore essential for predicting future forest dynamics. To address this gap, we performed a controlled lethal drought experiment with *Fagus sylvatica* seedlings, applying a population-level genome-wide association approach that regresses phenotypic class-mean drought survival against allele frequencies of the same classes from whole-genome SNP data, assuming an additive contribution of the trait loci (Pfenninger, 2025). Controlled experimental designs offer a pragmatic framework to resolve the ambiguities of multiple co-occurring stressors that are difficult to disentangle under field conditions; by isolating soil drought as a single, discrete stressor, we can map genomic regions to specific phenotypic traits and establish functional genotype–phenotype links. We monitored individual survival length as a direct proxy of fitness. Specifically, we aimed to: 1) resolve phenotypic variation in response to lethal drought conditions, 2) identify the genomic basis of this variation and 3) compare these results with selected sites from field data (Eberhardt, Reuss, Nieto Blázquez, et al., 2026). Altogether, our analyses aim to clarify whether standing genomic variation can explain differential drought survival in European beech seedlings, and to link these phenotypic manifestations to genotypic candidate loci with relevance for predicting forest regeneration success under future climate scenarios.

## 2. Material & Methods

### Seed preparation & cold stratification

Beech nuts were collected in autumn of 2023 (collection sites presented in S2). The beech nuts were sterilised in 5-10% NaOCl, washed 2x with deionised water and then dried superficially on paper towels. The beech nuts were stratified by storing at 2°C and -5°C for 4 weeks each. After thawing at 2°C for 24 hours, the nuts were watered for another 24 hours and finally stored at 3°C until germination. An additional 2000 beech nuts harvested in 2022 were ordered from a forest seed bank. The nuts had been frozen up until dispatch from the seed bank. Upon arrival at the institute, the nuts were immediately weighed, measured and planted upon germination. All plants were raised under controlled conditions in a laboratory at 20°C and 1000 Lux light intensity.Seedlings were grown in pots (0.6L) with drainage holes and watered with a fixed volume (50 mL per cell every second day) sufficient to bring the substrate to field capacity, with excess water draining freely.

### Experiment

Once a plant had two fully developed leaves, it was transferred to a botanical climate chamber for the drought experiment. The following traits were measured at the experiment start: plant height, stem diameter, canopy diameter, length and width of each leaf. Two 5mm disks were punched from the cotyledons and stored immediately at -80°C for subsequent pool-sequencing. The botanical chamber was set to the following conditions: 16:8 hours day and night conditions (3000:0 Lux, cold white light), respectively, constant humidity of 60% and temperature of 18°C. Upon experiment start, the seedlings were not watered anymore. Soil moisture was measured on a subset of plants (n = 113) using the TRIME-PICO64 TDR sensor (IMKO, Ettlingen, Germany) once per week, as sensor availability was limited during the experimental period. Despite the low frequency and coverage, weekly measurements were sufficient to confirm the expected progressive decline in soil moisture across the drought treatment. These measurements were used to confirm drought progression rather than as a predictor in the statistical models. (S3). Seedling death was defined as complete wilting of the primary leaves. Photosystem II performance was assessed weekly using the Pocket PEA chlorophyll fluorimeter (Hansatech, Norfolk, England), measuring two complementary indices of photoinhibition and stress response: the maximum quantum efficiency of photosystem II (Fv/Fm), which reflects the proportion of absorbed light energy used for photochemistry in dark-adapted leaves (minimum 20 minutes) and declines under drought-induced oxidative stress and photoinhibition, and the Performance Index (PI), a multiparametric indicator that integrates the density of active reaction centres, the efficiency of primary photochemistry, and the capacity for downstream electron transport, rendering it a particularly sensitive early indicator of drought stress even before visible symptoms appear. At the end of the experiment (i.e. plant death), the following traits were measured: plant height, stem diameter, canopy diameter, length and width of each leaf, and root length. The plant was dried for 24 hours at 60°C and subsequently the above- and below-ground biomass of the dried plants were weighed.

To assess the relationship between individual morphological and physiological traits and drought survival, we fitted a series of univariate hierarchical Bayesian linear regression models using the brms package (Bürkner 2017) in R, with one model per trait. Both the predictor and the response variable were standardised to zero mean and unit variance prior to modelling, so that posterior estimates reflect effect sizes in units of standard deviations and are directly comparable across traits. In each model, the number of days survived until death was included as the continuous response variable with a Gaussian likelihood, and the respective trait was included as a continuous fixed-effect predictor. All models showed satisfactory convergence (R□ < 1.01, Bulk ESS > 1000) and adequate posterior predictive fit. Posterior predictive checks revealed a bimodal distribution in survival length that was not fully captured by the Gaussian likelihood, likely reflecting systematic differences in survival between origin groups. We therefore additionally visualised survival distributions separately by origin to characterise this pattern. Seed origin (n = 2 populations) was included as a fixed effect to control for systematic differences in survival between populations. Given the small number of origin levels, a fixed-effects parameterisation was preferred over a random-effects approach.. All models were fitted using brms default weakly informative priors. We report posterior means and 95% credible intervals for all fixed effects; predictors whose 95% credible interval excluded zero were considered to have a credible association with drought survival.

### DNA extraction, construction of pools and sequencing

Genomic DNA was extracted from 12.5 mm² leaf disks using the NucleoMag Plant Kit (Macherey Nagel, Düren, Germany) and samples were grouped into 8 pools by length of survival. Sequencing libraries (350 bp insert size) were prepared by Novogene (Cambridge, UK) and sequenced on the Illumina NovaSeq 6000 platform (150 bp paired-end) to a target depth of 40X per pool. Raw sequence data are available via the European Nucleotide Archive (ENA) (Eberhardt, 2023) under accession PRJEB64934.

### Data processing & mapping

After initial quality control with FastQC (Andrews, 2010), reads were trimmed using Trimmomatic v0.3.9 (Bolger et al., 2014), merged with PEAR v.0.9.11 (Zhang et al., 2014), and aligned to the reference genome (Mishra et al., 2022) using BWA-MEM v.0.7.17 (Li & Durbin, 2013). Post-alignment processing, including sorting, merging, and duplicate removal, was performed using SAMtools v.1.10 (Danecek et al., 2021) and Picard v.2.20.8 (Broad Institute, 2019). Final quality metrics were generated using Qualimap v.2.2.1 (García-Alcalde et al., 2012) before converting alignments to mpileup format using SAMtools v.1.10 (Danecek et al., 2021). The toolbox Popoolation2 (Kofler et al., 2011) was used to convert the mpileup to sync file format and to remove indels. From the sync file, which stores raw read counts, a custom Python script was implemented to convert the data to allele frequencies relative to the reference allele. The script filters the dataset for quality and representativeness, retaining SNPs with a minimum depth of 15X per pool, a minor allele frequency (MAF) > 0.1, and < 25% missing data.

To identify genetic variants associated with drought resilience, a popGWAS (Pfenninger, 2025) was performed. This approach runs a linear regression between the eight pools and survival means, thereby testing whether the allele frequency at each variable site correlates with survival across pools, thus assuming an additive trait model. Significant associations were visualised via QQ and Manhattan plots using the R package qqman v. 0.1.9 (Turner, 2018) to assess whether p-values deviate from what would be expected under the null hypothesis.

### Allele frequency polarisation and phenotypic gradient analysis

To investigate the relationship between allele frequencies at candidate loci and drought tolerance phenotype, we first identified the top 0.05% of SNPs from the poolGWAS results (n = 2,384 SNPs) based on –log10(p) values, retaining one SNP per 1 kb window to reduce linkage non-independence (n = 2,156 SNPs after filtering). Drought tolerance phenotype was quantified as the first principal component (PC1) of a PCA on individual-level phenotypic data, which was then averaged per pool to obtain a single PC1 score per pool. Pool PC1 means ranged from –2.67 (pool P1, least drought-tolerant) to 4.50 (pool P8, most drought-tolerant), with pool P4 representing the gradient midpoint (PC1 mean = -0.28). Allele frequencies (AF) were polarised so that the respective allele was increasing in frequency along the drought tolerance gradient. Polarisation was performed by regressing AF across the eight pools against pool-level PC1 means, and flipping AFs for SNPs where the regression slope was negative. This regression-based approach was preferred over a simple P1-vs-P8 comparison as it uses information from all eight pools and is more robust to pool-level noise at individual loci. To quantify realised and remaining adaptive potential, we calculated three delta AF metrics per SNP: P8 - P4, the AF difference between the gradient midpoint and the maximum phenotype; P8 - P1, the phenotypic range across the full gradient; and 1 - P8, the remaining potential for allele frequency change until fixation. Distributions of these delta AF values were visualised as histograms and compared using a Bayesian ANOVA framework implemented in the R package brms (Bürkner, 2017). Given that delta AF values are bounded between 0 and 1 and the 1 - P8 distribution contained a substantial spike at zero (n = 395 SNPs already fixed in P8), we modelled delta AF using a zero-one-inflated Beta distribution. The reference group was P8 - P4. Model convergence was assessed via Rhat values and visual inspection of posterior trace plots. Directional hypotheses were tested using posterior probabilities and evidence ratios derived from 4,000 post-warmup draws across four chains.

### Analysis of candidate loci

To characterise the biological roles of the candidate loci, we identified genes associated with the top 0.05% SNPs by intersecting SNP coordinates with the *Fagus sylvatica* genome annotation (GFF3 file). A SNP was assigned to a gene if its position fell within the genomic range (start to end) of a gene feature defined in the annotation file, including non-translated 3’ and 5’ regions. Gene Ontology (GO) terms and functional domains were previously assigned to the beech genome with the InterProScan database (v. Jan. 2025) (Paysan-Lafosse et al., 2023). A GO enrichment analysis was performed on the top 0.05% SNPs for ‘Biological Processes’ using the R package topGO v.2.24.0. (Alexa & Rahnenführer, 2009). Overrepresented terms were identified using Fisher’s exact test with the ‘classic’ algorithm, applying a significance threshold of p<0.05.

### Comparison to field data

To see whether the selected loci from the drought experiment play a role in the selection observed in the field, we compared the selected alleles to publicly available SNPs from a Poolseq forest dataset between canopy trees (BA, established ca. 1910–1930) and young growth (JU, established ca. 2000-2020) collected from the warmest and driest sites in the federal state of Hesse, Germany (Eberhardt, Reuss, Nieto Blázquez, et al., 2026). Selected loci in the latter were identified using a Cochran-Mantel-Haenszel (CMH) test to detect consistent allele frequency shifts between growth classes across populations, resulting in 1.3 million loci. The top 0.05% SNPs of both datasets were matched to search for overlaps in exact SNPs, genes, 1kb and 2 kb windows. A GO enrichment analysis (TopGO) was performed on the overlapping SNPs. All supporting data and scripts are archived on Zenodo (Eberhardt, Reuss, & Pfenninger, 2026) 10.5281/zenodo.21334604.

## 3. Results

### Phenotypic traits

The survival length varied by up to 30 days between the seedlings. The first individuals already died after six days without water, while the toughest seedlings lasted 40 days (Fig. 1a). Survival length showed a bimodal distribution across all seedlings, with the two modes corresponding to the two origin groups: seed bank-origin seedlings survived markedly fewer days than nursery-origin seedlings (Fig. 1b). The randomly collected beech nuts from the forest lasted 31 days under drought conditions on average (95% highest density interval (HDI) = 29 – 33), while the commercial seeds died after 21 days on average (95% HDI = 20 – 22), supported by large confidence (95% HDI = 8-13 days, pp (β> 0) = 100%, effect size Cohen’s D = 1,6; Fig. 1c). Especially among the most drought resistant individuals there were almost none from the seed bank.

**Fig. 1.**
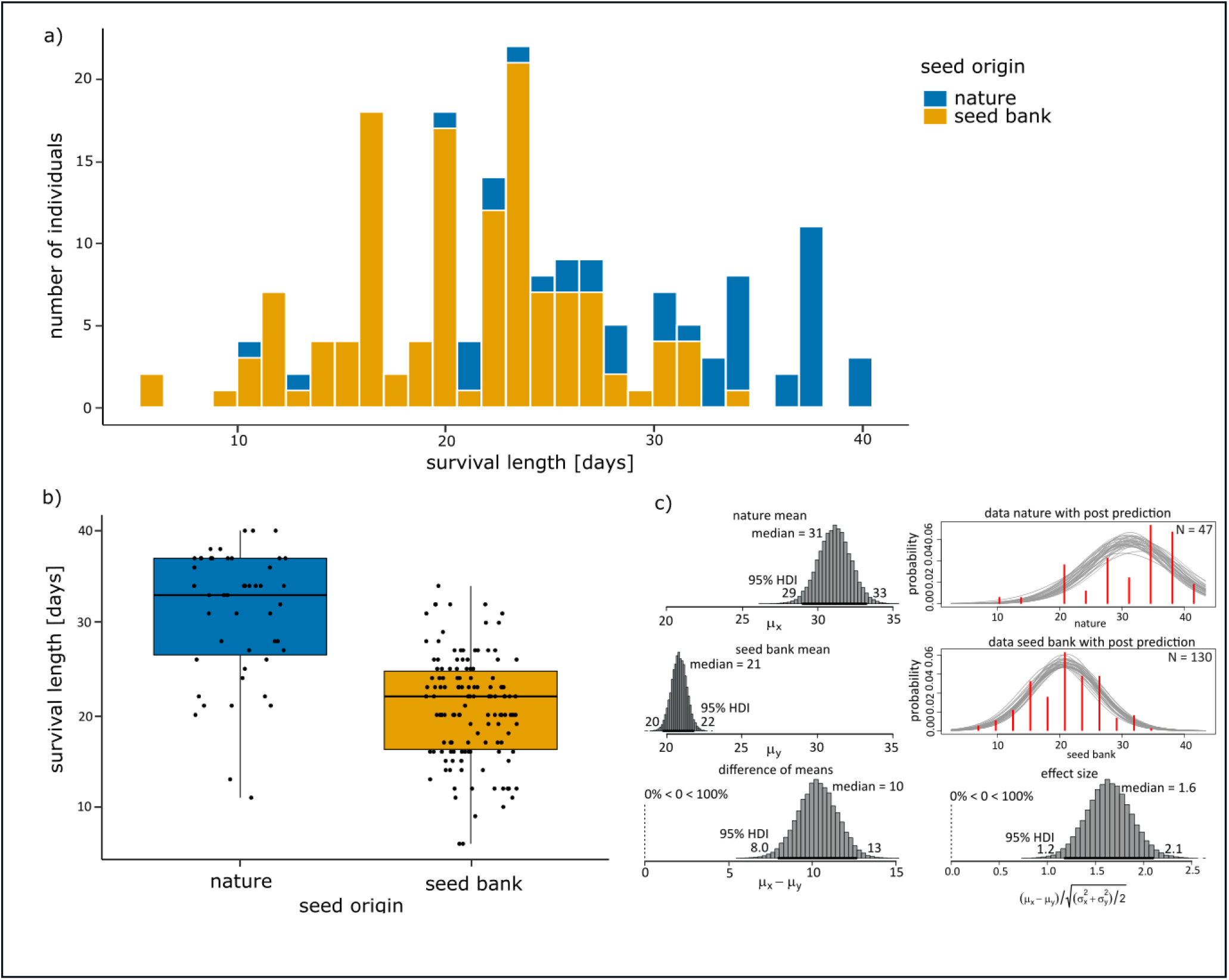
Distribution of survival length and influence of seed origin. a) Distribution of survival length by origin b) Survival length grouped by seed origin. c) Bayesian implementation of a t-test testing for a difference in survival length between collected seeds and seeds from a certified seedbank

This origin difference was reflected in the univariate hierarchical Bayesian linear regression models as a large negative fixed effect of seed bank origin (posterior mean = −0.96 SD, 95% CI [−1.24, −0.67]), regardless of the phenotypic predictor included in the model. Root traits, i.e. higher below-ground biomass, root:shoot ratio and longer roots (Fig. 2) were strongly positively correlated with survival length (β = 0.45 SD, 95% CI [0.33, 0.58]; β = −1.01 SD, 95% CI [−1.28, −0.75]; β = 0.32 SD, 95% CI [0.21, 0.43]). respectively), which was also evident in the PCA (S11). However, there was only a linear correlation within the first 30 days, and there was no positive effect of the trait on survival length after 40 days. Above-ground traits, i.e., above-ground dry mass and leaf area were strongly correlated with survival (β = 0.40 SD; 95% CI [0.06, 0.28]); β = 0.26 SD, 95% CI [0.14,0.38], respectively.) All results of the Bayesian linear regression models are presented in S4.

**Fig. 2.**
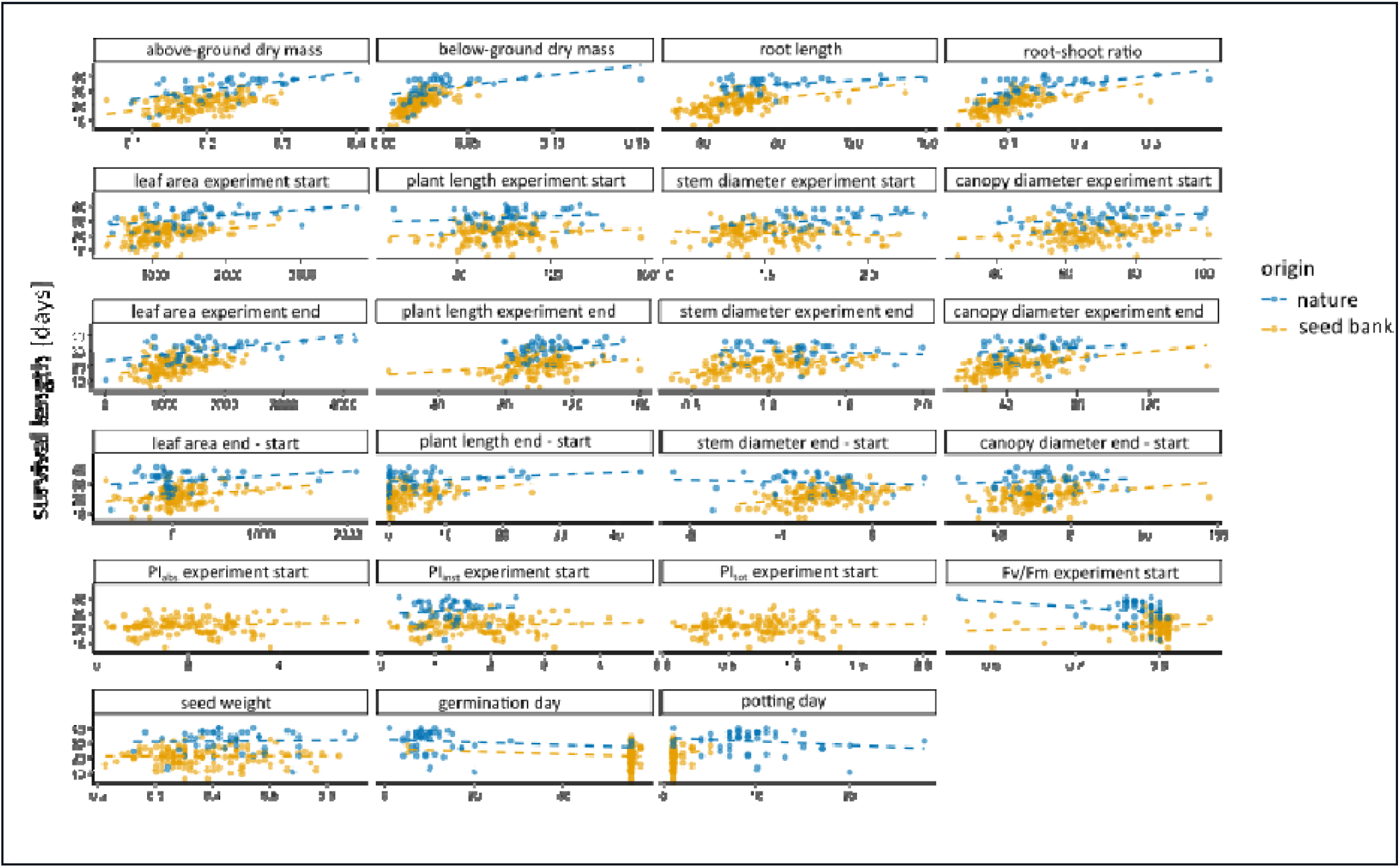
Simple biplot of measured traits against survival length grouped by seed origin.

### Genomic analysis

A Principal Component Analysis (PCA) on all measured traits showed that survival length best explained the data (PC1: 30,3 %; PC2: 14,1 % variance) (S11). We used this trait to group the individuals into eight clusters, from shortest to longest survival under the drought treatment. After quality filtering, pools had a mean coverage of 16X. From an initial 21 million positions, we identified 4.4 million SNPs. The popGWAS performed on pool-sequencing data from the seedling drought experiment identified 4.6 million SNPs from an initial 21 million positions, with a genomic inflation factor demonstrating a fitted slope of 0.84, indicating a degree of genomic deflation rather than inflation. While this suggests that population stratification was strictly controlled, the downward deviation of the observed test statistics relative to the null expectation indicates a conservative model fit or potential over-correction (S6). Plotting the mean polarised AF of these SNPs against the PC1 scores based on the measured phenotypic traits showed a consistent trend across the top candidate loci, indicating a strong support for the GWAS signal being biologically meaningful (Fig. 3a). The top 0.05% of significant loci were extracted for further analysis, resulting in 2,385 SNPs.

**Fig. 3.**
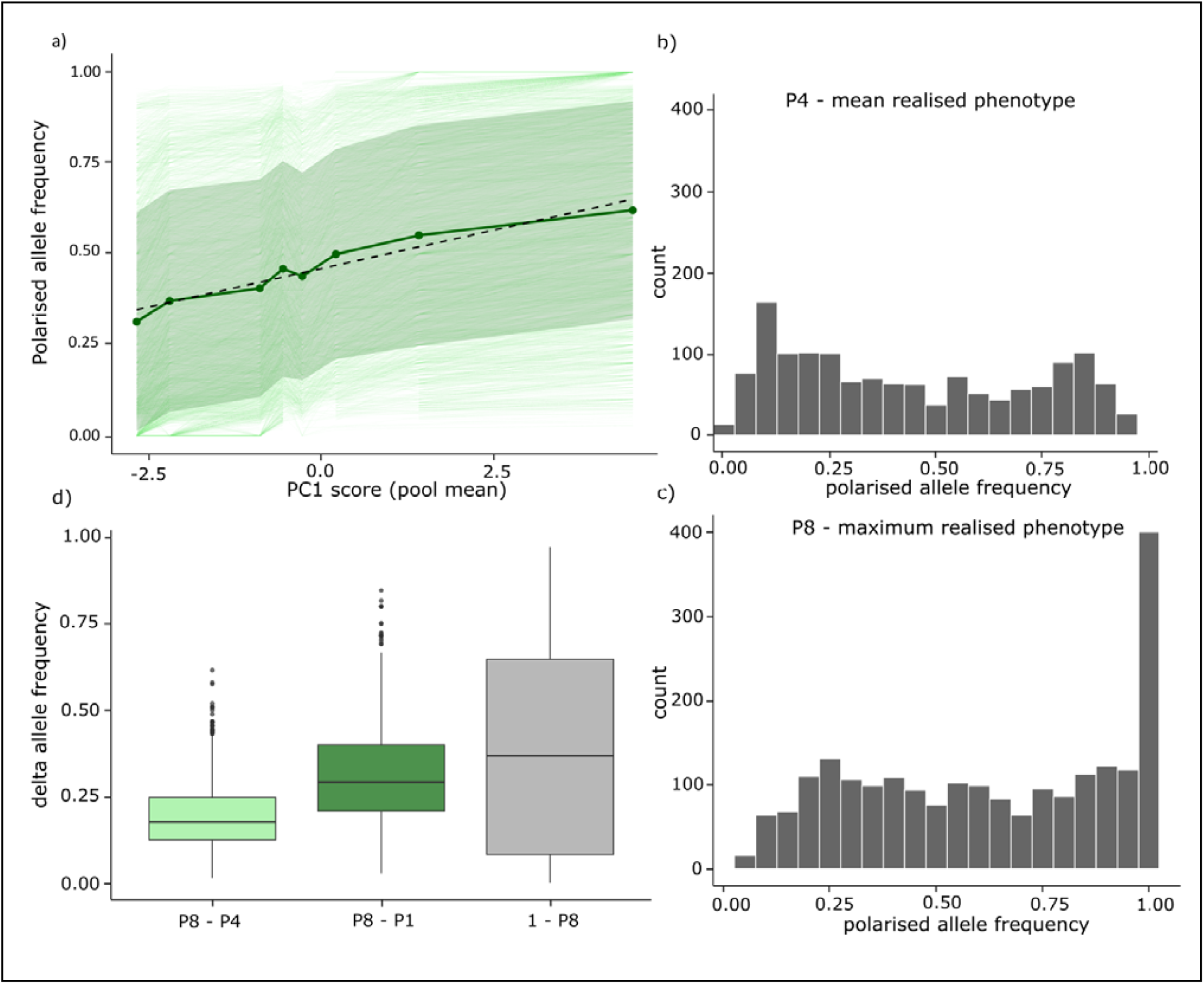
a) Phenotypic pool means expressed as PC1 scores plotted against allele frequencies at significant loci, polarised by increasing drought resilience. The solid green line connects the mean allele frequencies per class over all significant loci, the hashed line the calculated regression line. The shaded area denotes the 95% confidence interval b) Allele frequency distribution of candidate loci in the phenotype pool showing average drought resilience (P4) c) Allele frequency distribution of candidate loci in the phenotype pool showing maximum drought resilience (P8) d) Boxplot of allele frequency differences between maximum and mean phenotype (left), maximum and minimum phenotype (middle), and remaining potential for allele frequency changes until fixation (right)

### Allele frequency increases along the drought tolerance gradient

After regression-based polarisation, mean AFs of candidate loci increased monotonically from P1 to P8 across the pool PC1 gradient, confirming that polarisation successfully oriented all alleles in the direction of increasing drought tolerance (Fig. 3a). The AF distributions at the gradient midpoint (P4) and maximum (P8) differed markedly: At P4, allele frequencies showed a bimodal distribution with peaks at low (AF ∼0.10) and high (AF ∼0.75–0.85) frequencies (Fig. 3b), while at P8 the distribution was strongly right-skewed with a pronounced spike at AF = 1 (Fig. 3c), indicating that a substantial proportion of candidate loci are already near or at fixation in the most drought-adapted pool.

### Realised and remaining adaptive potential

The total realised AF range across the gradient (P8 - P1) was centred around 0.25-0.30, indicating that already a modest but consistent directional increase in AFs can confer the observed phenotypic differences (Fig. 3d). The mean AF difference from the gradient midpoint to the maximum (P8 - P4) was accordingly narrower and centred around 0.15- 0.20, reflecting the smaller phenotypic range covered by the upper half of the gradient (Fig. 3d). The remaining adaptive potential (1 - P8) showed a large range, driven by a right-skewed distribution with the highest frequency at zero, indicating that many loci are already fixed in the most drought-tolerant pool, while a substantial proportion (∼ a third) retain moderate-to-large remaining potential for a further increase in AF (Fig. 3d). Bayesian ANOVA confirmed that all three delta AF groups differed decisively from one another (Fig. 4d). The direct comparison between 1 - P8 and P8 - P1 confirmed that the mean remaining adaptive potential per locus exceeds the mean realised allele frequency difference across the full gradient (posterior mean difference = 0.55, 95% CI [0.51, 0.59] (S12). All three hypotheses had evidence ratios of infinity and posterior probabilities of 1, indicating that not a single posterior sample contradicted the stated directional effects. Model convergence was excellent, with all Rhat values equal to 1.00 and well-mixed posterior trace plots across all four chains (S12). The posterior predictive check indicated that the zero-one-inflated Beta model captured the overall shape of the data reasonably well, though the bimodal structure of the combined delta AF distribution was only partially reproduced, reflecting the known limitation of a single shared shape parameter across groups.

### GO enrichment analysis with topGO

GO enrichment analysis of the most significant 0.05% SNPs revealed significant overrepresentation of genes involved in transcription and RNA processing, carbohydrate metabolism and protein glycosylation, translation and protein targeting, chromosome organisation and meiosis, cellular signalling and enzyme regulation, metabolism and biosynthetic processes (S8).

### Partial overlap of involved loci with field data

The top 0.05% SNPs between old and young growth, BA and JU, showed enriched gene functions related to vesicle transport and exocytosis, cellular localization and transport, RNA processing and ribosome biogenesis, chromosome organisation and genome architecture, metabolic process and modification and biological regulation (S9).

The 0.05% top SNPs of the two datasets shared 24 SNPs within 2kb windows. These were enriched for functions related to plant development and morphogenesis, cellular, root and epidermis development, kinase activity and phosphorylation signalling, metabolic and catalytic regulation, and lipid transport and homeostasis (S10). Of those shared alleles, four were found within 1kb of those SNPs that showed large selection coefficients in Q4, the warmest quartile, one being an exact match, although only one of the SNPs was annotated (Bhaga_10.g1558) (S7). Based on the functional annotation from the Valley Oak Genome Project, gene Bhaga_10.g1558 (NCBI Reference Sequence: XP_030939709.1) is predicted to encode a receptor-like protein kinase (RLK) and shows sequence homology to the Lr10 disease-resistance locus, which is associated with rust fungus resistance (Sork et al., 2016).

## 4. Discussion

The results of this study provide insight into the physiological and genomic basis of drought tolerance in *Fagus sylvatica* seedlings, with consistent signals emerging from both phenotypic trait associations and population-level genomic analyses. Strong variation in phenotypic drought response among seedlings were mirrored in the functional annotation of candidate loci, indicating considerable potential of young growth to adapt to changing environmental conditions as well as current phenological mismatch (Fig. 4b).

### Phenotypic Predictors of Drought Survival

The strong positive association between root traits and survival length is consistent with the well-established importance of below-ground resource acquisition under drought stress (Guo et al., 2024; Kou et al., 2022; Shoaib et al., 2022). Higher below-ground biomass and longer roots likely confer a competitive advantage under soil water deficit by enabling seedlings to access deeper or wider soil moisture reserves, a strategy documented in drought-tolerant tree species, such as Douglas fir, Scots pine and limber pine (Lazarus et al., 2018; Moser et al., 2016). The positive effect of root:shoot ratio on survival further supports this interpretation. Cork oak seedlings were shown to allocate proportionally more biomass to roots relative to shoots, thereby buffering against hydraulic failure during drought, as the root system can sustain water uptake while shoot demand remains comparatively low (Morillas et al., 2023).

Above-ground traits, while also significantly associated with survival, showed weaker effect sizes compared to root traits, suggesting they play a secondary role in drought tolerance in this species at the seedling stage. The association between above-ground biomass and leaf area with survival may reflect the general vigour of seedlings rather than specific drought-adaptive strategies. Larger, more vigorous seedlings may simply have greater resource reserves to draw upon during drought, a finding consistent with positive effects of root mass, root length and total mass on seedling survival during a natural drought (Walters et al., 2023). Nevertheless, these traits should not be dismissed, as leaf area in particular influences transpiration rates and water loss, and its positive association with survival may seem counterintuitive. This may indicate that seedlings with greater initial leaf area had already undergone stomatal regulation.

The correlation between planting day and survival length is likely attributed to the fact that seedlings from the seed bank germinated later than the collected beech nuts. Interestingly, seed weight did not correlate with drought resistance. This contradicts literature linking heavier seeds to higher lipid content and superior drought tolerance through transgenerational effects (Hatzig et al., 2018; Herman et al., 2012; Kalandyk et al., 2017). If the heavier beech nuts could be traced back to drought stress in the parental generation causing higher lipid content, we would have expected a correlation of seed weight and survival length in the experiment. However we did not have information on the cause of the observed variation in seed weight which may reflect factors irrelevant for seedling fitness under drought stress. Moreover, the studies on transgenerational effects focused on crop plants, which possess less standing genetic diversity which could enable plastic responses to drought; in such systems, larger individuals generally withstand stress better. In contrast, environmental stress in highly diverse forest trees may unveil adaptive plasticity driven by cryptic genetic variation (Pfenninger et al., 2025).

### Origin as a Predictor of Survival

The large and consistent effect of seed origin on survival length was an intriguing finding. To understand the contrasting performance between the seed bank-derived seedlings and the wild-collected cohorts under drought, potential confounding factors such as seed storage age must first be considered. Seeds from the seed bank were harvested one year prior to the self-collected seeds and stored frozen. Storage can incur physiological costs that impair seedling vitality and stress tolerance, independent of genetic background (Ratajczak et al., 2015). However, such an effect was observed only after at least three or more years (Walters et al., 2023; Wawrzyniak et al., 2020) and has therefore likely not played a decisive role here. According to the literature, seedlings from certified sources sometimes show comparable (Buiteveld et al., 2025), but generally much better, growth performance than randomly collected seeds (Dedefo et al., 2017; Grossnickle & Ivetic, 2017). However, these comparisons were conducted under optimal growing conditions. Our explanation for the observed phenomenon is that the trees from which the seeds were obtained were themselves selected for optimal growth under optimal conditions from a forestry perspective, a practice that was sensible under the stable climatic conditions of the last 250 years. Under the experimental drought conditions, however, they perform worse than the beechnuts collected in nature, which additionally originate from a large number of genetically variable trees rather than a limited number of selected seed trees. This effect could be even greater for seedlings raised in nurseries. The extent of this effect requires further investigation, as it has direct implications for restoration and reforestation practices: If material from seed banks is systematically less drought-tolerant, then the currently applicable legal requirements regarding seed origin could be actively undermining efforts to establish climate resistant forests under the increasingly arid conditions predicted for Central Europe.

### Evolutionary potential

The monotonic increase in mean allele frequencies across all eight pools, rather than only between phenotypic extremes, provides empirical support for the validity of the incremental pooling approach implemented in the poolGWAS framework of (Pfenninger, 2025). Unlike conventional pool-seq GWAS designs that contrast the most extreme phenotypic pools, this approach assigns pools along a continuous phenotypic gradient, thereby preserving quantitative variation in AF trajectories. The recovery of a consistent, gradient-wide directional signal at candidate loci suggests that the method successfully captures the polygenic architecture of drought tolerance rather than artefacts of extreme-pool contrasts, such as population structure or environment-specific effects unrelated to the trait of interest (Pfenninger, 2025).

The pronounced right-skew in the P8 AF distribution, with a spike at AF = 1, reflects that the most drought-resistant individuals in this experiment share a high proportion of candidate alleles at or near fixation (Fig. 3c). Rather than indicating directional selection in the classical sense, this pattern suggests that the extreme end of the phenotypic distribution is characterised by individuals that happen to carry the adapted allele at high frequency across many candidate loci, consistent with a polygenic basis of drought tolerance where phenotypic extremes arise from the cumulative effect of many loci each contributing a small effect (Pritchard & Di Rienzo 2010). This pattern additionally reflects the allelic composition and diversity present at the sites where the beech nuts were sampled. The bimodal AF distribution at the gradient midpoint (P4), with peaks at low (∼0.10) and high (∼0.75–0.85) frequencies and a dip at intermediate values (∼0.50), likely reflects the genetic structure of the experimental material (Fig. 3b). As the beech nuts for this experiment were sourced from two sites, the candidate loci may segregate at characteristically low or high frequencies depending on the allelic composition of each source population, resulting in a bimodal rather than uniform AF distribution across the gradient. The dip at intermediate values (∼0.50) is expected, as loci with no consistent directional trend across the gradient cannot rank among top GWAS candidates. The delta AF analyses reveal a wide phenotypic range with a median realised range of ∼0.25–0.30 across the full phenotypic gradient (Fig. 3d). The remaining adaptive potential is larger than the currently realised range. This indicates that the candidate loci identified here collectively harbour considerable scope for further allele frequency change under intensified drought stress, suggesting substantial adaptive capacity of European beech under climate change scenarios (Fig. 3d).

### Genomic Basis of Seedling Drought Tolerance

Our GO enrichment analysis revealed a significant enrichment of RNA splicing pathways in drought-resilient individuals, highlighting post-transcriptional regulation as a crucial mechanism in the drought response of *Fagus sylvatica* seedlings. Alternative splicing (AS) often serves as a rapid mechanism that stabilises photosynthetic complexes, enhances reactive oxygen species (ROS) scavenging, and limits oxidative damage under water deficits (Collin et al., 2025; Grasser, 2025; Huang et al., 2024). The enrichment of mRNA splicing implies that fine-tuning isoform expression patterns is a core adaptive strategy for coping with severe drought (Filichkin et al., 2018; Xu et al., 2023).

Concurrently, the enrichment of glycosylation-related pathways underscores the role of post-translational modifications in driving abiotic stress tolerance. Protein glycosylation ensures structural stability under environmental stress and protects membrane and cell-wall proteins from dehydration-induced denaturation (Chang et al., 2021). This modification participates in post-translational crosstalk: for example, the N-glycosylation of cell-wall-associated Receptor-Like Kinases (RLKs) is required for proper folding and ligand perception (Nekrasov et al., 2009). The genomic variations observed in both glycosylation enzymes and kinase domains suggest a coordinated optimisation of this sensing apparatus, through which phosphorylation signals under stress are transferred downstream (Gandhi & Oelmüller, 2023). Beyond structural protection, glycosyltransferase activity regulates abscisic acid (ABA)-independent pathways that amplify stomatal closure and the accumulation of protective osmolytes, like proline and flavonoids (Verma et al., 2026). Drought-induced stomatal closure also confirms the previous considerations regarding the measured larger leaf area in drought-resistant seedlings.

The enrichment of carbohydrate and polysaccharide biosynthetic processes points to a dual strategy of cellular protection and energy management. Both structural and soluble polysaccharides optimise water-storage capacity under severe water deficits and maintain cell wall integrity against osmotic shock (Ezquer et al., 2020; Zhao et al., 2025). Pathways for sucrose, starch, and galactose metabolism are co-regulated with antioxidant biosynthesis, thereby offering both osmotic protection and redox control (Haghpanah et al., 2024). These metabolic processes are reinforced by significant genomic variation within NAD/H metabolic pathways, which preserve Photosystem II function and combine with ABA signalling networks to adjust downstream stress-responsive genes (Hong et al., 2020; Mittler & Jones, 2023). Furthermore, enriched GTPase regions highlight an evolutionary advantage that optimises stomatal water conservation while maintaining resistance against fungal pathogens (Ganotra et al., 2022).

The enrichment of chromosome organisation terms suggests that processes involving the genome structure contribute to drought tolerance through multiple mechanisms. Methylation and chromatin remodelling improve accessibility of stress-response genes and preserve genome integrity under water deficit (Nguyen et al., 2022; Sow et al., 2021). Gene dosage effects through tandem duplication may further amplify drought-protective functions, as shown for dehydrin genes in *Caragana* where copy number enhanced drought survival (Mei et al., 2025). Additionally, variation in regulatory architecture, such as promoter and haplotype variation, can shift transcriptional programmes governing xylem development, ABA signalling, and growth-defence trade-offs (Fang et al., 2023).

Finally, the enrichment of genes involved in meiotic recombination addresses a critical gap in forest climate-adaptation research. While individual-level drought stress can mechanistically induce higher meiotic crossover frequencies to generate novel allelic combinations (Verde et al., 2025), wild populations exposed to long-term climate stress may exhibit reduced effective recombination rates (Eberhardt, Reuss, Nieto Blázquez, et al., 2026). This discrepancy likely reflects the difference between the immediate cellular response and long-term ecological outcomes. While stress may trigger meiotic hyper-recombination at the individual level to generate novel allelic combinations, intense multifactorial field stress drives severe demographic bottlenecks and selective sweeps. These in turn increase the proportion of homozygous sites in the genome and thus the possibility to detect recombination (Hartfield & Bataillon, 2020; Peñalba & Wolf, 2020). In natural stands, strong selective pressure preserves intact, co-adapted gene complexes in surviving cohorts, ultimately resulting in the genome-wide signature of reduced effective recombination.

Ultimately, these insights demonstrate that drought tolerance in *Fagus sylvatica* is a highly polygenic trait rooted in fundamental cellular maintenance. Seedling survival length under extreme physiological stress seems to be mainly dictated by baseline genetic efficiency in RNA processing, protein folding, energy homeostasis, and macromolecular repair.

### Shared SNPs

The overlapping SNPs between the wild *Fagus sylvatica* climate-adaptation dataset (Eberhardt, Reuss, Nieto Blázquez, et al., 2026) and the controlled seedling drought experiment converge into three functional clusters: Leaf and root morphology, cellular development, stress signalling and homeostasis. SNPs associated with signalling and homeostasis are significantly enriched for lipid localisation and transport terms. This functional suite serves as a central hub for drought adaptation by integrating the reinforcement of physical barriers, such as the cuticle, with cell membrane homeostasis and hormonal stress signalling (Cameron et al., 2006; Liang et al., 2023). For instance, the overexpression of lipid transfer proteins has been shown to enhance early seedling growth and root elongation in *Arabidopsis* (Jan et al., 2026). Given that overall vigour and a robust root architecture were associated with prolonged survival in the seedling experiments, lipid transport likely represents a critical genetic mechanism linking root development to systemic physiological drought tolerance.

Furthermore, we identified enriched terms nested under the regulation of transferase activity, specifically targeting protein serine/threonine kinases, cyclin-dependent kinases, and phosphorylation cascades. As previously discussed, these phosphorylation networks are essential components of upstream stress-sensing pathways. Crucially, phosphorus metabolism acts as a metabolic core during drought stress; it directly modulates ABA signalling to control stomatal closure, fuels rapid phospholipid-mediated signalling cascades, and facilitates cellular energy triage under deficit. This shared genomic signature suggests that climatic selection targets these upstream osmotic signalling cascades, which are responsible for translating turgor loss into adaptive cellular responses.

The concurrent enrichment of lipid signalling, kinase activity, and membrane homeostasis GO terms among the shared SNPs implies that natural selection has acted on a dual front: the capacity to rapidly perceive osmotic shock at the membrane level and the ability to sustain cellular function during prolonged water deficits. This aligns with a dual strategy of stress anticipation and tolerance. These cellular mechanisms translate directly into the structural adjustments seen in mature forests. Changes in the plant epidermis and root system morphology demonstrate how mature trees adapt to water deficits through foliar modifications, such as altered stomatal size and increased cuticle thickness, alongside shifts in root architecture that optimise water uptake and storage. Furthermore, these wild-tree adaptations provide strong evidence that the phenotypic advantages observed in our seedling trials, specifically an increased root:shoot ratio, persist into maturity under natural forest conditions.

Lastly, while GO terms related to multicellular organismal processes are inherently broad, they are driven by a distinct set of loci separate from those governing leaf and root morphology. These genes likely govern fine-scale cell size, stomatal development, or xylem differentiation, which collectively minimise transpirational water loss and prevent lethal xylem embolism. Ultimately, changes to root, leaf, and cellular development represent long-term structural adjustments to macro-climatic shifts. Conversely, the pathways involved in stress signalling and cellular homeostasis enable rapid, plastic responses to fluctuating environmental conditions. This dual architectural and physiological rewiring highlights an evolutionary survival strategy that facilitates phenotypic adjustments to changes in both the climatic mean and its standard deviation. *Fagus sylvatica* has experienced oscillating climatic changes associated with the Pleistocene (Kremer et al., 2025), the observed selection may thus represent active adaptive tracking to large-timescale environmental changes (Pfenninger & Foucault, 2022).

### Adaptive potential to mismatch among seedlings

The drought experiment revealed substantial phenotypic and standing genetic variation among seedlings. The genomic signature extracted from the seedling drought experiment reflects immediate, short-term survival mechanisms under lethal water deficits, relying heavily on osmoprotectant accumulation, strict stomatal regulation, and rapid stress-signalling cascades. Intriguingly, not a single enriched GO term could be directly linked to macro-scale root morphological development. Rather than indicating a lack of structural adaptation, this suggests that the observed drought-resistant phenotypes are driven by a complex, coordinated interplay of alternative splicing, flexible signalling networks, and starch-to-sugar metabolic biosynthesis.

The identified candidate loci from the drought experiment revealed high phenotypic variability to climate-change related mismatches in young growth. Despite the near fixation of a substantial number of candidate loci in the most drought-adapted pool, an immense adaptive potential remains in the population that even exceeds the realised range to date. Assuming that this potential for increase in allele frequencies in this polygenic trait translates in respective phenotypes, these findings suggest that European beech retains substantial genomic capacity for further adaptation to drought stress, with important implications for its resilience under projected climate change scenarios.

Comparison of candidate loci identified here with those detected in field populations under natural conditions (Eberhardt, Reuss, Nieto Blázquez, et al., 2026). revealed that the experimentally identified loci represent a subset of the broader selective signal detected in the wild. This overlap is notable given the fundamental differences between the two datasets: the present study identified candidate loci under controlled drought conditions in young seedlings, isolating drought stress as the primary selective pressure, whereas field-sampled trees are older individuals that have been exposed to the cumulative effects of multiple, co-occurring selective forces including heat, drought, radiation, and pathogen pressure over decades. The substantial overlap between these two datasets therefore not only provides partial validation of the controlled experiment, but also suggests that drought tolerance is a primary and persistent driver of natural selection in wild beech populations, with the controlled experiment successfully capturing a core subset of the genomic variation under selection in the field.

The consistent and large effect of seed bank origin across all analyses highlights the importance of seed provenance in drought tolerance studies and should be accounted for in both experimental design and restoration planning. Moreover, it shows that performance under stress cannot be inferred to performance under optimal conditions. Given the large-scale planting efforts following drought-induced forest die-off, this finding should be verified urgently to prevent further maladaptation in the next generation of trees.

## Acknowledgments

We thank Lothar Volk from HessenForst for the water content measurements of our seeds and his help with the germination, and the Molecular Ecology group for their help with the beech nut collection.

## Statements and Declarations

The authors declare no conflict of interest.

## Data availability statement

The data that support the findings of this study are openly available. Raw sequence data are deposited in the European Nucleotide Archive (ENA) under accession PRJEB64934 (Eberhardt, 2023). Supporting data and scripts are archived on Zenodo (Eberhardt et al., 2026) 10.5281/zenodo.21334604).

